# Stress limits the beneficial effects of glutamine in male *ob/ob* mice

**DOI:** 10.64898/2026.07.21.739777

**Authors:** Adam Tiffay, Candice Lefebvre, Charles-Edward Breemeersch, Virginie Dreux, Christine Bôle-Feysot, Charlène Guérin, Elise Maximin, Magali Monnoye, Pierre Déchelotte, Véronique Douard, Alexis Goichon, Moïse Coëffier

## Abstract

**Introduction:** Obesity is a major health issue associated with metabolic and psychological comorbidities, as well as an increased prevalence of disorders of gut–brain interaction (DGBI). Obesity and DGBI share common mechanisms such as inflammation, gut barrier dysfunction, and alterations of gut microbiota, which are all known to be regulated by stress. Glutamine (Gln), which is essential to maintain intestinal integrity and immune response, may counteract these alterations. This study aimed to evaluate the effects of oral Gln supplementation on stress-induced response in obese mice.

**Methods:** Seven-week-old male leptin-deficient *ob/ob* mice were assigned to four groups: control, chronic restraint stress (CRS), Gln-supplemented, or both CRS and Gln-supplemented. Gln was administered in drinking water for two weeks, and CRS was performed during the final 4 days. Metabolic parameters, intestinal permeability, inflammatory markers, gene and protein expression, and gut microbiota composition were assessed.

**Results:** Stress increased plasma corticosterone levels but had a limited effect on metabolic parameters. In obese mice without stress, Gln supplementation reduced body weight gain, improved body composition and reduced inflammation in the visceral adipose tissue. These effects were lost under stress conditions, with an increase in fasting glycaemia. Stress reduced occludin protein levels, while Gln exerted context-dependent effects, decreasing gene expression of *Tjp3, Cldn15* and *Ccl2* in unstressed mice but increasing gene expression of multiple tight junction (*Tjp2*, *Tjp3*, *Cldn12*, *Cgn*, *F11r*, *Marveld2*) and inflammatory markers (*Tlr2*, *Myd88*, *Irf3*) under stress. Interestingly, in unstressed obese mice, Gln altered the composition of the gut microbiota, with changes in key bacterial taxa (*Thermodesulfobacteriota* and *Clostridiaceae*). This was associated with decreased levels of cecal short-chain fatty acids and increased levels of branched-chain fatty acids.

**Conclusion:** In conclusion, Gln improves metabolic and adipose inflammatory parameters in genetically obese mice. However, these benefits are no longer observed when mice are under stress conditions. Since, Gln has been found to increase fasting glycaemia and colonic inflammation, in association with alterations of gut microbiota.

## Introduction

Obesity has become a major public health concern in both industrialized and developing countries, with more than 50% of the European population classified as overweight or obese (1). For the first time, in 2025, the prevalence of obesity surpassed that of underweight in school-aged children and adolescents aged of 5–19 years according to a UNICEF report (2). Obesity, that is defined by a body-mass index (BMI) value of 30 kg/m² or greater, is a complex multifactorial disease resulting from an interaction between genetic predispositions, metabolic dysregulations and environmental factors. Individuals with obesity present an increased risk of developing multiple comorbidities, including metabolic syndrome, dyslipidemia (3), depressive disorders, coronary artery diseases and type 2 diabetes mellitus (4). Recently, a higher prevalence of disorders of the gut-brain interaction (DGBI) in patients with class III obesity in the European adult population has been reported (5). Obesity and DGBI share common pathophysiological mechanisms, including low-grade systemic inflammation, intestinal microbiota dysbiosis and an impaired intestinal barrier (6,7). Stress contributes greatly to the onset, maintenance and exacerbation of symptoms in patients with DBGI (8). Nevertheless, the links that explain the relationships between psychological stress and gastrointestinal symptoms during obesity remains still unknown.

Several studies suggested that psychological stress increases secretion of corticotropin-releasing factor (CRF), leading to mast cell activation and the release of mediators that disrupt mucosal barrier function (9–11). Water avoidance stress (WAS), commonly used to mimic Irritable Bowel Syndrome (IBS) symptoms in rodent, exacerbated intestinal and colonic hyperpermeability, respectively, in obese mice fed with high-fat diet (HFD) and deficient for leptin (*ob/ob* mice) (12). In the chronic-restraint stress (CRS), increased serotonin levels, exacerbated oxidative stress, gut dysbiosis and disrupted colonic barrier function have been shown in non-obese mice (13,14). In male obese mice, CRS was able to affect body weight gain, glucose homeostasis and cecal microbiota (15,16). Considering these data, new strategies targeting both gastrointestinal symptoms and metabolic disorders may be of interest during obesity.

Glutamine (Gln) is a non-essential amino acid and preferential substrate for rapidly dividing cells. Previous research in multiple animals models has demonstrated that Gln enhances the proliferation of intestinal epithelial cells, stem cells, and crypt cells, thereby facilitating the repair of injured intestinal mucosa (17–20). Moreover, Gln affects the composition of gut microbiota, strengthens intestinal innate immunity, and mitigates excessive inflammatory cytokine production (21–23). Interestingly, oral Gln supplementation was able to reduce abdominal pain and normalize intestinal hyperpermeability in IBS patients with diarrhea (24). In animal models of stress with gastrointestinal symptoms, Gln prevented colonic hyperpermeability in mice subjected to WAS (25) or CRS (26). Gln also appears as a good candidate to limit obesity-associated intestinal and metabolic disorders in rodents (16,27) and humans (28,29).

Therefore, the present study aimed to evaluate the impact of oral Gln supplementation on CRS response in a genetic-model of obesity, the leptin-deficient *ob/ob* mice, by assessing gut barrier function, inflammatory responses, gut microbiota, and metabolic complications.

## Material and methods

### Animal experimentation

Seven-week-old male *ob/ob* mice (B6.V-*Lep^ob^*/JRj, n=40, n=4/cage) were obtained from Janvier Labs (Le Genest-Saint-Isle, France) and acclimatized for 4 days in a controlled environment (23°C with a 12 hours light-dark cycle) with free access to water and a Standard Diet (SD, 3.34 kcal/g, Cat # 1314, Altromin). Throughout the protocol, from week 1 (W1) to week 9 (W9), mice were fed with SD. Mice were randomized into four groups: (i) *ob/ob*-NS group (mice without Gln supplementation neither restraint stress), (ii) *ob/ob*-S group (mice with CRS), (iii) *ob/ob*+Gln group (mice with Gln supplementation), and (iv) *ob/ob*-S+Gln group (mice with Gln supplementation and CRS). Body weight was measured weekly, and body composition was measured by EchoMRI (EchoMRI, Houston, US) on W7 and W9. At the end of the protocol, the mice were anesthetized by injection of a mixture of ketamine (100 mg/kg; Cat # KTP07, Boehringer Ingelheim) and xylazine (10 mg/kg; Cat# 047-956, Rompun, Bayer Healthcare) followed by cervical dislocation. Blood samples were collected, centrifuged at 3000*xg*, 4°C for 15 min and plasma was frozen at −80°C. Cecal content, colon, subcutaneous (scWAT) and perigonadal (pWAT) adipose tissues were collected and immediately stored at −80°C until analysis. The protocol was approved by the regional ethics committee (authorization on APAFIS #29283-2021012114574889 v5) and performed in accordance with the current French and European regulations.

### Oral Gln supplementation

From W7 to W9, mice from *ob/ob*+Gln and *ob/ob*-S+Gln groups received an oral Gln supplementation in drinking water to provide 2 g.kg^−1^ of body weight each day. This Gln concentration corresponds to a dose at 0.16 g/kg /day in humans (24) and has already used in chronic-stress animal models (16,26). The Gln solution was prepared freshly and replaced every day.

### Chronic restraint stress

At W9, mice from *ob/ob*-S and *ob/ob*-S+Gln groups were subjected to CRS for 2 hours per day during the last 4 days of the protocol. Briefly, mice underwent a brief isoflurane anesthesia (5%, <3 min) and were subsequently confined in restraint cages (Bioseb®, Vitrolles, France) for two hours before being released back into their home cages. They were subjected to the same restraint sessions for four consecutive days at the same hour. This model of stress has been validated by Langlois *et al.* and control mice remained in their home cages throughout the procedure (26).

### Oral glucose tolerance test (OGTT)

An oral glucose tolerance test (OGTT) was conducted before (W7) and after Gln supplementation (W9). Following a 12 hour overnight fasting period with unrestricted access to water (without Gln), mice received an oral dose of glucose (1 g/kg of body weight; Cat# G8769, Merck) by gavage. Blood glucose levels were then measured in the caudal vein at intervals of 0, 15, 30, 45, 60, and 90 min using a glucometer (Cat# 4015630082841, Accu Check Guide, Roche).

### In vivo and ex-vivo intestinal permeability

I*n vivo* intestinal permeability was evaluated by measuring the appearance of FITC-dextran in the blood following oral administration as previously described (30). Briefly, following a 2-hour fasting period and three hours before euthanasia, the mice received a FITC-dextran solution (4 kDa, 40 mg/mL; Cat # FD4-1G, Merck) by oral gavage at a dosage of 10 µL per gram of body weight. The fluorescence intensities of FITC-dextran (λ excitation at 485 nm, λ emission at 535 nm) in plasma were measured with the Spark multimode microplate reader (Tecan, UK).

Colonic paracellular permeability was also evaluated by measuring fluxes of Alexa Fluor 680-dextran (3 kDa, 0.02 mg/mL; Cat # D34681, Merck) and Lucifer Yellow (400 Da, 2.5 mg/mL, Cat# L453, Merck) across colonic mucosa mounting in Ussing Chambers (Cat# MA1 66-00XX, Harvard Apparatus) after 3 hours at 37°C. Fluorescence intensities were measured using the Spark multimode microplate reader (Tecan, Life Sciences) with following parameters: Alexa Fluor 680-Dextran (λ excitation at 665 nm, λ emission at 710 nm) and Lucifer Yellow (λ excitation at 428 nm, λ emission at 540 nm) in the serosal medium.

### Plasma assays: obesity and inflammation markers

Plasma levels of insulin (Cat# 171G7006M), leptin (Cat# 171G7007M), resistin (Cat# 171G7009M), adiponectin (Cat# 171F7002M), IL-6 (Cat# 12002241), and TNFα (Cat# 12002444) were measured through Bio-Plex Pro assays (BioRad Laboratories) according to the manufacturer’s guidelines, employing Luminex technology (BioRad Laboratories). Additionally, plasma concentrations of urea (Cat# 98-11070-01), alanine aminotransferase (ALAT, Cat# 98-11067-01), aspartate aminotransferase (ASAT, Cat# 98-11069-01), and total cholesterol (Cat# 98-11072-01) were determined using a Catalyst One analyzer (IDEXX Laboratories).

### ELISA - Quantification of Plasma Corticosterone

Plasma levels of corticosterone were measured using the Corticosterone ELISA kit (Cat# KA0468; Abnova Corporation) according to the manufacturer’s guidelines.

### Colonic RNA extraction and RT-qPCR

Total RNA was isolated from colonic mucosa using the TRIzol protocol (Cat# 15596018, Invitrogen, Fisher Scientific) after mechanical homogenization. RNA concentration was determined with a NanoDrop 2000 spectrophotometer (Cat# ND-2000, Fisher Scientific). One microgram of RNA was then reverse transcribed into cDNA using an M-MLV reverse transcriptase in the presence of random hexamers and RNase inhibitor as previously described (30). Quantitative PCR analyses were performed on a QuantStudio 12K Flex real-time PCR system (Life Technologies) using a Master Mix Fast SYBR™ Green (Cat# 4385612, Life Technologies). The panel included 44 target genes related to inflammatory pathways and intestinal barrier function as previously used (30). The detailed primer sequences are provided in this previous publication. Gene expression was normalized to the geometric mean of *Actb* and *Gapdh*.

### Adipose tissue RNA extraction and RT-qPCR

Total RNA from perigonadal (pWAT) and subcutaneous (scWAT) adipose tissues was extracted using the RNeasy Lipid Tissue Mini Kit (Cat# 74804, QIAGEN) according to the manufacturer’s instructions. RNA concentration and reverse transcription were performed as described for colonic tissue. Quantitative PCR was then performed using SYBR Green chemistry on a Bio-Rad CFX96 real-time PCR system (Bio-Rad Laboratories) targeting selected cytokines (*Il6*, *Tnfa*, *Ccl2*), adipokines (*Lep*, *Retn*, *Adipoq*) and Gln transporter *Slc1a5* (F: CATGTAAAATACCGCAATCCTGT; R: GACGATAGCGAAG ACCACCA, Tm: 62°C). Expression levels were normalized to the mean of *B2m* and *Rps18*. Primer sequences are detailed in a previous article (30).

### Protein extraction and Western blotting

Colon samples were homogenized using the Tissue Lyser LT (Cat# 85600, QIAGEN) with buffer A (50 mM HEPES pH 7.5, 150 mM NaCl, 10 mM EDTA, 10 mM glycerophosphate, 100 mM NaF) or U-CHAPS lysis buffer before subjected to Western blotting(31,32). The extraction of proteins from pWAT and scWAT tissues was performed using the RNeasy® Lipid kit (Cat# 74804, QIAGEN), in which the organic phase containing proteins. After the addition of isopropanol, washing steps with guanidinium chloride (0.3 M in 95% ethanol), the samples were rinsed with 100% ethanol and reconstituted in 200 µL of 2X Laemmli buffer with 5% β-mercaptoethanol (v/v). The protein content in the supernatants was quantified using Bradford assay or the 2D Quant kit (Cat# GE80-6483-56, Cytiva) for proteins extracted with the U-CHAPS buffer (31,32). 25 µg of total proteins were subjected to denaturation at 95°C for 5 min in a solution containing 2X of Laemmli buffer and 5 % (v/v) of β-mercaptoethanol (Cat# M131, Merck). Subsequently, the protein samples were loaded into the wells of a 4-20% Mini-PROTEAN TGX gels (Cat# 4568096 or Cat# 4568093, Bio-Rad Laboratories) and underwent electrophoresis for 1.5 hours at 125V (40 mA/gel). The proteins were then transferred to a PVDF (Cat# 1620177, Bio-Rad Laboratories) or nitrocellulose membrane (Cat# 10600008, Cytiva) using a liquid-phase electro-transfer system from BioRad. The membranes were incubated for 1 hour in a blocking solution consisting of 5% skim milk (w/v) or 5% BSA (w/v) and TBS-T [0.5 M Tris, 0.05% Tween 20 (v/v)]. The primary antibody was incubated overnight at 4 °C, followed by three 5-min washes in TBS-T and a 1-hour incubation with the anti-mouse (Cat# P0161, Agilent; 1/5000) or anti-rabbit (Cat# P0399, Agilent; 1/5000) secondary antibody at room temperature. Five primary antibodies were used in the present study: anti-occludin (Cat# 33-1500, ThermoFisher Scientific, 1/1000; Nitrocellulose; 5% skim milk), anti-ZO-1 (Cat# 40-2200, ThermoFisher Scientific, 1/2000; PVDF; 5% skim milk), anti-claudin 1 (Cat# 71-7800, ThermoFisher Scientific, 1/1000; PVDF; 5% BSA), anti-O-Linked N-Acetyl-glucosamine (O-GlcNAc, Cat# ab273; Abcam, 1/2500; PVFD; 5% skim milk). Protein bands were visualized using the Clarity™ Western ECL Substrate kit (BioRad) and the ChemiDoc™ XRS+ system (BioRad Laboratories), followed by quantification and visualization using Image Lab 6.0.1 software (BioRad Laboratories). The expression of each protein of interest was normalized with total protein expression levels using the “Stain-free imaging” mode.

### Cecal microbiota: DNA extraction, sequencing, and data processing

Fresh cecal samples were collected from the diapers and stored at −80°C until microbiota analysis. Total DNA was extracted using DNA extraction QIAamp PowerFecal Pro kit (QIAGEN). The V3-V4 hypervariable region of the 16S rRNA gene (33) was amplified using Phanta Max Super-Fidelity DNA Polymerase (Vazyme) and the primers forward V3F: 5′-CCTACGGGNGGCWGCAG-3′ and reverse V4R: 5′-GACTACHVGGGTATCTAATCC-3’ were used (95°C for 3 min and then 35 cycles at 95°C for 15 sec, 65°C for 15 sec, and 72°C for 1 min before a final step at 72°C for 5 min). The primers were selected from the Klindworth *et al.* publication (34). The purified amplicons were sequenced using Miseq sequencing technology (Illumina) at the @BRIDGe sequencing facility (GABI, INRAE, AgroParisTech, Paris-Saclay University, Jouy-en-Josas, France).

The resulting sequences were analysed using R (Team, 2019) workflow combining DADA2v.1.36 (35) and FROGS 5.0.0 (36) software on the Migale Bioinformatics server (Université Paris-Saclay, INRAE, MaIAGE, 78350, Jouy-en-Josas, France). Reads were processed using DADA2, including filtering with filterAndTrim and ASV inference using the core sample inference algorithm. Chimeric sequences were removed using VSEARCH. ASVs with a relative abundance below 0.005% were filtered out. Taxonomic assignment was performed using the FROGS affiliation tool against the NCBI 16S rRNA reference database (v1.20230726) as previously described (30).

### SCFAs analysis

Short-chain fatty acids (SCFAs) were extracted from cecal contents, precipitated with phosphotungstic acid, and analyzed by gas chromatography using 2-ethylbutyrate as internal standard, as previously described (30).

### Statistical analysis

Data were analyzed with GraphPad Prism 8.0.2 (GraphPad Software Inc., San Diego, CA, US) and expressed as the mean ± standard error of the mean (SEM). Body weight and glycaemia were compared using a two-way ANOVA (time × groups), followed by Tukey’s post hoc tests. For other parameters, a two-way ANOVA (Gln × Stress) was performed with Tukey’s post hoc tests. Outlier detection was performed using an iterative Grubbs’ test at α = 0.05. For all analyses, statistical significance was set at p < 0.05, while a trend was considered for p-values between 0.1 and 0.05.

Bioinformatic analysis of 16S rRNA amplicon sequencing data was conducted in R (v4.4.2) using the Phyloseq package (v1.50.0). Alpha diversity index (Observed species, Chao1, Shannon and InvSimpson), phylum and family relative abundances were analyzed using a two-way ANOVA test followed by Bonferroni post-tests. Differential abundance analysis was performed using the DESeq2 package (v1.46.0).

## Results

### Effects of stress and Gln supplementation on body weight, glycaemia control and obesity-related plasma biomarkers

CRS induced an increase in plasma corticosterone levels [p(Stress)=0.0021], with a trend toward an increase in *ob/ob*-S mice compared to *ob/ob*-NS mice (p=0.0603, **Fig. 1A**). While CRS did not significantly affect body weight, body composition (**Fig. 1B-E**) and fasting glycaemia (**Fig. 1F**), a decrease in plasma insulin levels and HOMA-IR index was observed in *ob/ob*-S mice compared to *ob/ob*-NS mice (**Fig. 1G-H**). Apart from resistin levels, no other significant differences were observed in plasma parameters (**Fig. Supp. 1A-G**).

**Fig. 1.**
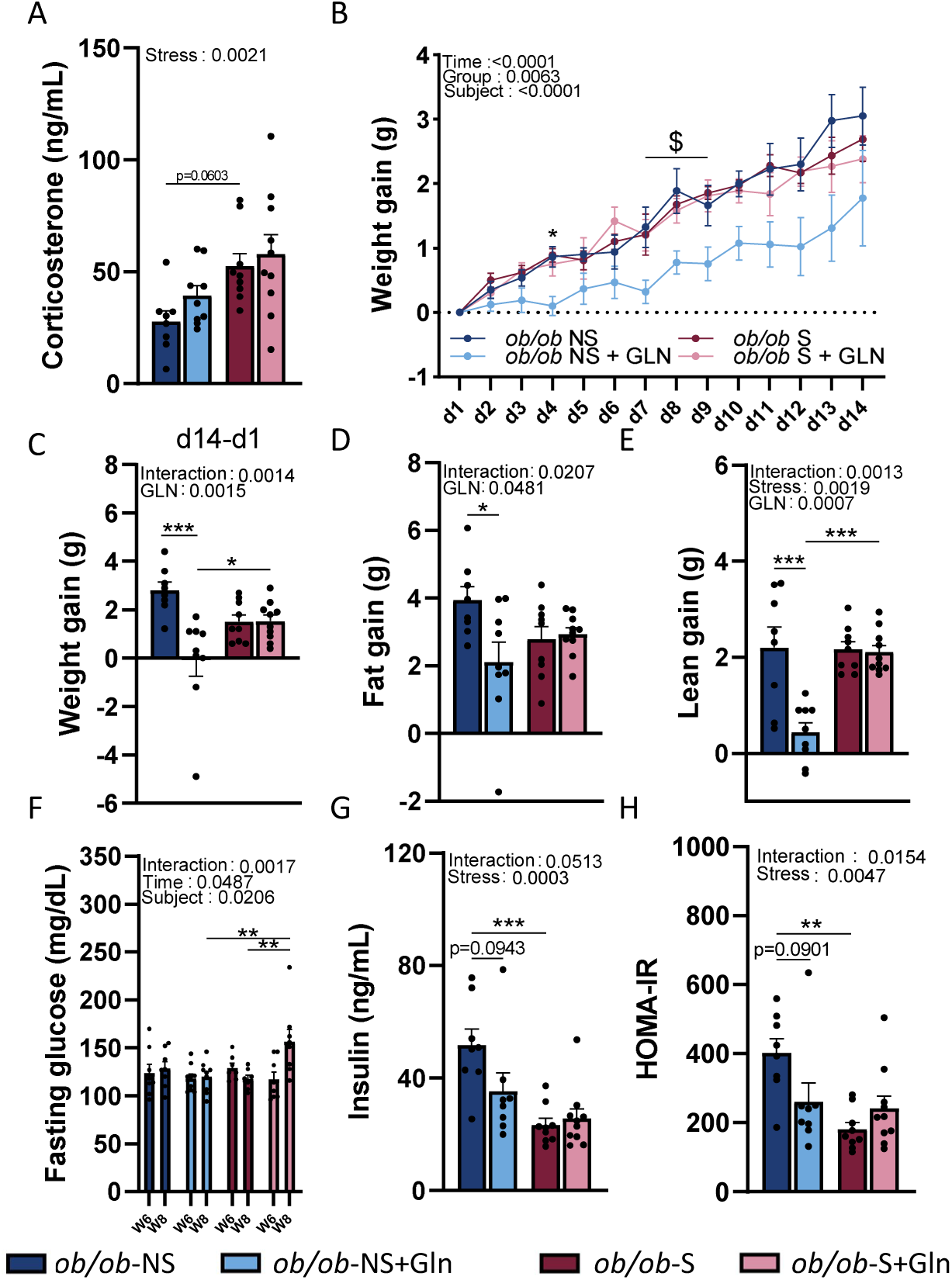
Plasma corticosterone, body composition and glycaemia parameters in male *ob/ob* mice supplemented or not with oral Gln and subjected to a CRS. Plasma corticosterone (ng/mL) **(A)**, kinetic weight gain (g) expressed as the difference between all day with the day before Gln supplementation **(B)**, delta weight gain (g) **(C)**, fat gain (g) **(D)** and lean gain (g) **(E)** (both measured by EchoMRI) expressed as the difference between the day before sacrifice with the day before Gln supplementation (d1-d14, 2g/kg body weight by day). Fasting glucose (mg/dL) **(F)**, plasma insulin (pg/mL) **(G)**, and HOMA-IR **(H)** expressed as [Fasting glucose (mmol/l) × insulin (mlU/L)/ 22,5]. Groups were compared with 2-way ANOVA followed by Tukey post hoc tests (*p < 0.05, **p < 0.01, ***p < 0.001). In (**B**), *, p < 0.05 *ob/ob*-NS vs *ob/ob*-NS+GLN and $, p < 0.05 *ob/ob*-NS+GLN vs *ob/ob*-S+GLN. Data are presented as mean ± standard error of the mean (SEM). n=9-10/group.

In unstressed *ob/ob* mice, Gln supplementation significantly reduced the gain in body weight (−2.89 g, p<0.001), fat (−1.83 g, p<0.05) and lean mass (−1.76 g, p<0.001) (**Fig. 1B-E**) but did not affect fasting glycaemia (**Fig. 1F**). A trend for a decrease in fasting plasma insulin levels and HOMA-IR index (−42%) was also observed but differences did not reach significance (**Fig. 1G-H**). Under Gln supplementation, CRS did not significantly increase corticosterone levels (**Fig. 1A**). By contrast, in stressed *ob/ob* mice, oral Gln failed to affect body weight and composition (**Fig. 1B-E**), insulin levels and HOMA-IR index (**Fig. 1G-H**) but increased fasting glycaemia (**Fig. 1F**, p<0.01). Gln supplementation and stress, each alone, did not modify plasma biomarkers related to obesity, except for plasma resistin level that was decreased by stress (**Table 1**). We also observed that plasma total cholesterol (p=0.0492) and ALAT (p=0.0031) were significantly increased in *ob/ob*-S+Gln mice compared to *ob/ob*-NS+Gln mice (**Table 1**). Taken together, these data suggested that (i) Gln supplementation reduces body weight gain and improves body composition in unstressed *ob/ob* mice, and (ii) these beneficial effects disappeared in stress conditions.

**Tab. 1.** Biochemical, metabolic, adipokine, incretin and inflammatory markers.

| Biochemical and Metabolic Markers |  |  |  |  |  |  |  |  |  |  |  |  |  |  |  |  |
| --- | --- | --- | --- | --- | --- | --- | --- | --- | --- | --- | --- | --- | --- | --- | --- | --- |
|  | Total Cholesterol (g/L) |  |  |  | Triglyceride (g/L) |  |  |  | ASAT (U/L) |  |  |  | ALAT (U/L) |  |  |  |
| Group | Mean | SEM | Significance Level | n | Mean | SEM | Significance Level | n | Mean | SEM | Significance Level | n | Mean | SEM | Significance Level | n |
| *ob/ob* -NS | 1,345 | ± 0.080 | a,b | 8 | 0,970 | ± 0.048 | ns | 8 | 459,125 | ± 76.381 | ns | 8 | 400,875 | ± 79.422 | a,b | 8 |
| *ob/ob* +Gln | 1,187 | ± 0.074 | a | 9 | 0,952 | ± 0.034 | ns | 9 | 450,111 | ± 69.074 | ns | 9 | 263,778 | ± 20.192 | a | 9 |
| *ob/ob* -S | 1,362 | ± 0.055 | a,b | 9 | 0,937 | ± 0.037 | ns | 9 | 390,333 | ± 46.010 | ns | 9 | 347,750 | ± 48.474 | a,b | 8 |
| *ob/ob* -S+Gln | 1,425 | ± 0.046 | b | 10 | 0,876 | ± 0.043 | ns | 10 | 587,000 | ± 34.586 | ns | 10 | 506,700 | ± 28.893 | b | 10 |
|  | GTT (U/L) |  |  |  | Urea (g/L) |  |  |  | Total protein (g/L) |  |  |  |  |  |  |  |
| Group | Mean | SEM | Significance Level | n | Mean | SEM | Significance Level | n | Mean | SEM | Significance Level | n |  |  |  |  |
| *ob/ob* -NS | 9,125 | ± 1.394 | ns | 8 | 0,436 | ± 0.036 | ns | 8 | 60,500 | ± 2.528 | ns | 8 |  |  |  |  |
| *ob/ob* +Gln | 8,222 | ± 1.103 | ns | 9 | 0,462 | ± 0.099 | ns | 9 | 56,111 | ± 1.703 | ns | 9 |  |  |  |  |
| *ob/ob* -S | 6,778 | ± 0.778 | ns | 9 | 0,274 | ± 0.005 | ns | 9 | 58,556 | ± 0.784 | ns | 9 |  |  |  |  |
| *ob/ob* -S+Gln | 6,500 | ± 1.167 | ns | 10 | 0,284 | ± 0.019 | ns | 10 | 60,500 | ± 1.621 | ns | 10 |  |  |  |  |
| Adipokine, incretin and Inflammatory Biomarkers |  |  |  |  |  |  |  |  |  |  |  |  |  |  |  |  |
|  | Adiponectin (pg/ml) |  |  |  | Resistin (pg/ml) |  |  |  | GLP-1 (pg/ml) |  |  |  | IL6 ( pg/ml) |  |  |  |
| Group | Mean | SEM | Significance Level | n | Mean | SEM | Significance Level | n | Mean | SEM | Significance Level | n | Mean | SEM | Significance Level | n |
| *ob/ob* -NS | 24490,686 | ± 2620.474 | ns | 8 | 273,280 | ± 22.896 | a | 8 | 0,433 | ± 0.053 | ns | 8 | 0,081 | ± 0.021 | ns | 7 |
| *ob/ob* +Gln | 29755,310 | ± 1825.669 | ns | 9 | 302,590 | ± 18.721 | a | 9 | 0,402 | ± 0.049 | ns | 9 | 0,097 | ± 0.028 | ns | 7 |
| *ob/ob* -S | 29001,126 | ± 1572.151 | ns | 9 | 208,783 | ± 12.273 | b | 9 | 0,332 | ± 0.027 | ns | 9 | 0,038 | ± 0.006 | ns | 8 |
| *ob/ob* -S+Gln | 27208,258 | ± 2368.645 | ns | 10 | 176,810 | ± 8.392 | b | 10 | 0,360 | ± 0.030 | ns | 10 | 0,070 | ± 0.021 | ns | 8 |

Plasma total cholesterol (g/L), triglycerides (g/L), ASAT (U/L), ALAT (U/L), GTT (U/L), Urea
(g/L), Total protein(g/L), Adiponectin (pg/ml), Resistin (pg/ml), GLP-1 (pg/ml) and IL6
(pg/ml) in male *ob/ob* mice (n=7-10/group). Groups were compared with 2-way ANOVA
followed by Tukey post hoc tests (Group with no common letter are consider as significative).
Data are presented as mean ± standard error of the mean (SEM).

### Effects of Stress and Gln supplementation on inflammation of white adipose tissue (WAT)

Perigonadal (pWAT) and subcutaneous (scWAT) white adipose tissues were analyzed by western blotting and RT-qPCR to determine the effects of Gln supplementation or stress on inflammatory component. In unstressed *ob/ob* mice, oral Gln reduced by 19% the protein expression of O-GlcNAcylation (**Fig. 2A-B**) and tended to decrease mRNA levels of *Tnfa* (**Fig. 2C**) and *Ccl2* (**Fig. 2D**) in pWAT, that were not observed in scWAT (**Fig. 2F-I**). These data indicated the beneficial effects of Gln on inflammation in pWAT in unstressed *ob/ob* mice. The mRNA levels of *Slc1a5*, which codes for a Gln uptake transporter, were reduced in pWAT of *ob/ob*-NS mice after Gln supplementation (**Fig. 2E**), but not in scWAT (**Fig. 2J**). Conversely, no significant effects of stress (CRS) or interaction (Stress × Gln) were observed in pWAT and scWAT. In scWAT, a “Gln” ANOVA effect was observed for mRNA levels of *Adipoq* [p(GLN)=0.0276] whereas no difference was detected in pWAT (**Fig. Supp. 1H-I**) and no difference was detected for *Il6*, *Il1b, Retn, Glul* and *Gls* in both pWAT and scWAT (**Fig. Supp. 2A, 2J**). These data highlighted the tissue-dependent effects of Gln on WAT inflammation.

**Fig. 2.**
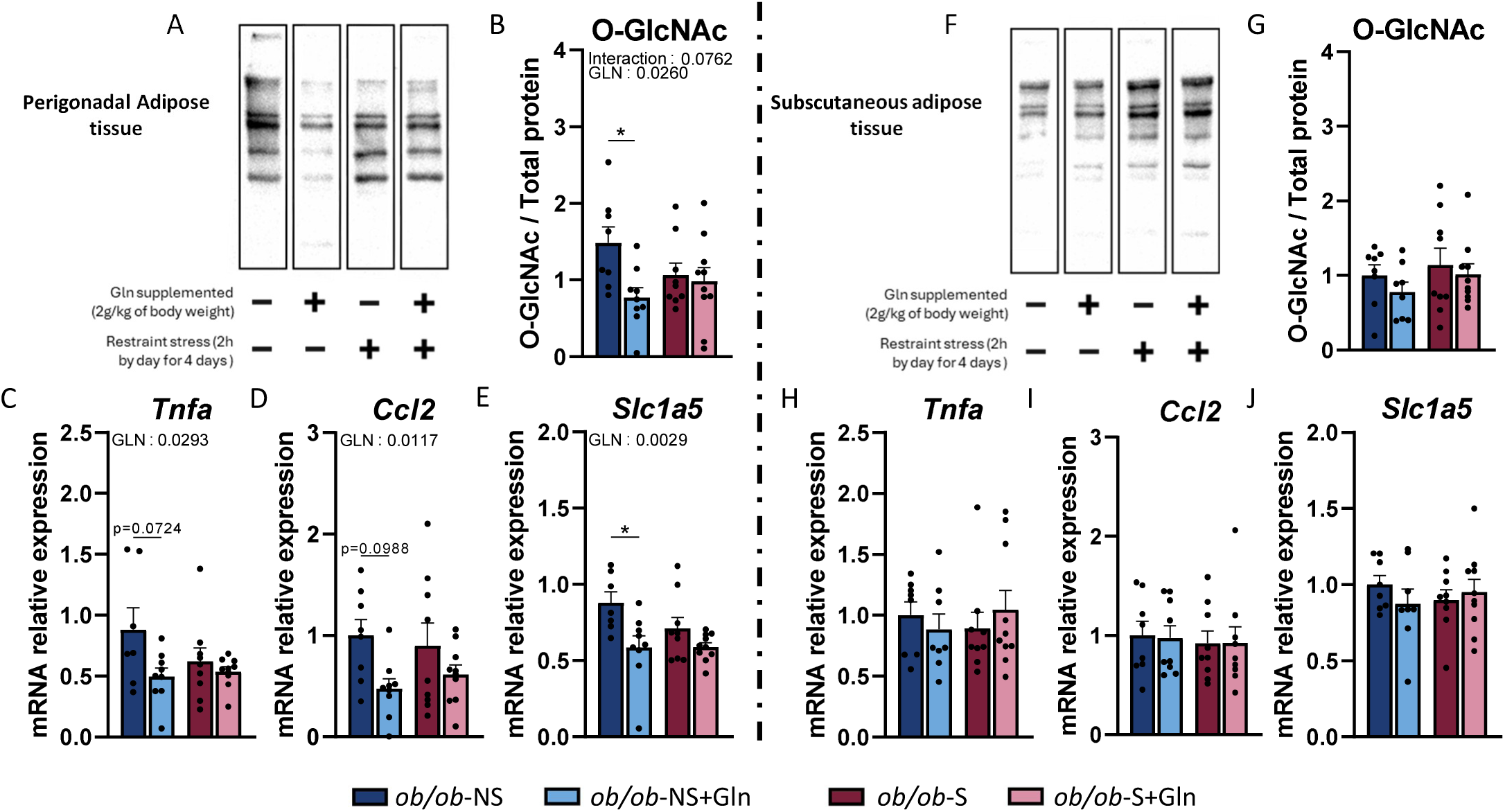
Inflammatory markers in perigonadal (A, B, C, D, E) and subcutaneous (F, G, H, I, J) adipose tissues of male *ob/ob* mice supplemented or not with oral Gln and subjected to a CRS. Cropped immunoblot membranes **(A, F)** and relative expression of O-GlcNac proteins **(B, G)** in adipose tissues of male *ob/ob* mice (n=9-10/group). Relative quantification of mRNA transcript levels inflammatory cytokine Tumor Necrosis Factor (*Tnfa*) **(C, H)**, chemokine ligand 2 (*Ccl2*) **(D, I)** and Neutral amino acid transporter B(0) (*Slc1a5*) **(E, J)**. Groups were compared with 2-way ANOVA followed by Tukey post hoc tests (*p < 0.05). Data are presented as mean ± standard error of the mean (SEM).

### Effects of stress and Gln supplementation on gut barrier and colonic inflammation markers

We examined gut permeability in mice and the colonic expression of 11 tights junction proteins and inflammatory markers by using western blotting analysis or RT-qPCR. However, neither stress nor Gln altered colonic paracellular and gut permeability (**Fig. 3A-C**). Stress reduced the protein expression of occludin (**Fig. 3D**), but not the *Ocln* mRNA levels (**Fig. 3E**), decreased the mRNA levels of *Cldn4* [p(stress)=0.0369, **Fig. 3F**] but enhanced the mRNA levels of *Tjp1* [p(stress)=0.0125, **Fig. 3G**] in the colon of *ob/ob*-mice. In unstressed *ob/ob* mice, Gln reduced mRNA levels of *Tjp3, Cldn15* [p(GLN)=0.0241, **Fig. 3H,I**] and *Ccl2* (**Fig. 4F**) in colonic tissue, whereas mRNA levels of *Fabp2* was increased (**Fig. 3J**). In stressed *ob/ob* mice, Gln supplementation enhanced the colonic mRNA levels of 5 tight junction proteins (*Tjp2*, *Cldn12, Cgn, F11r* and *Marveld2*; **Fig. 3 K-O**).

**Fig. 3.**
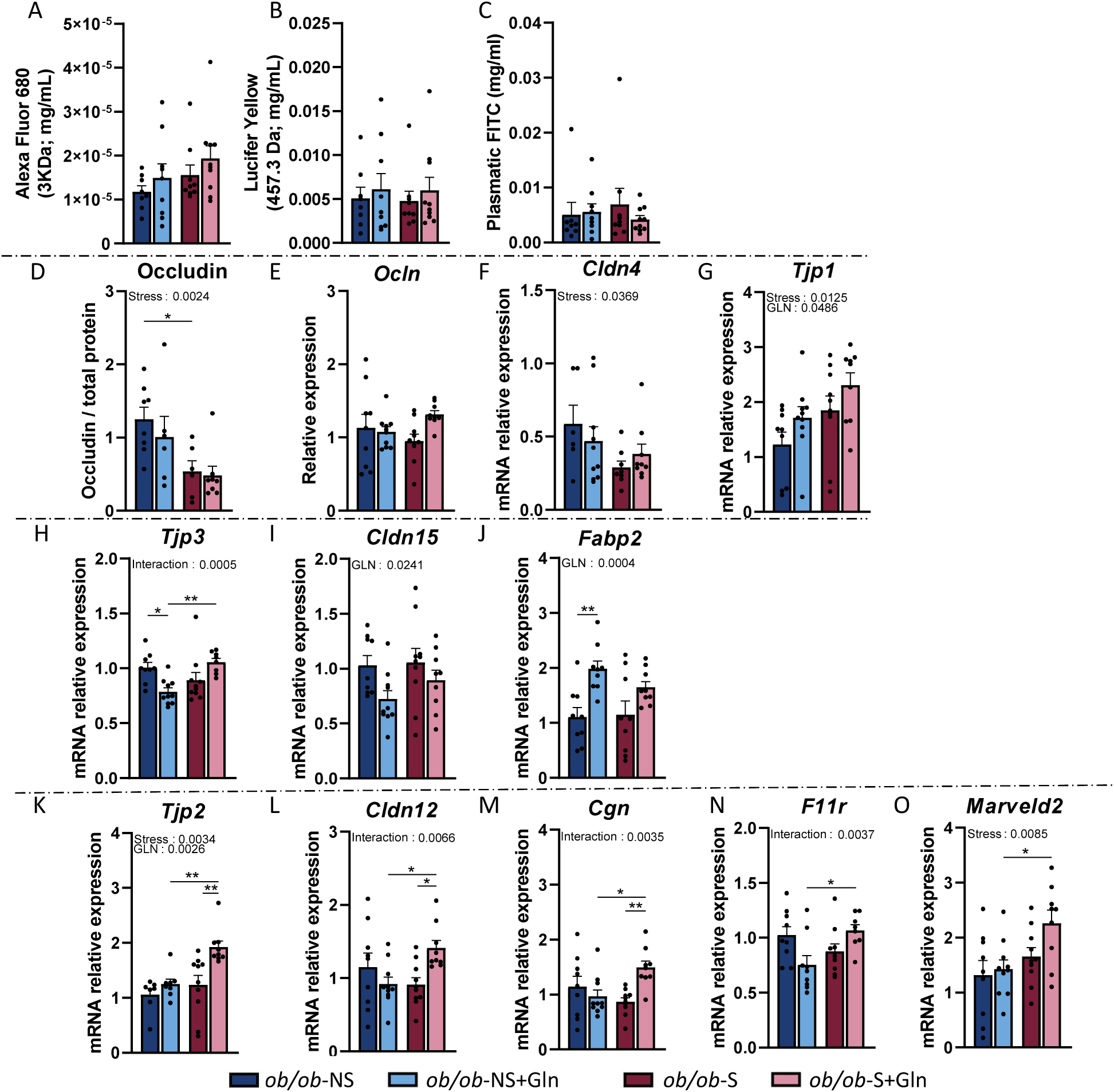
Intestinal Permeability and tight junction protein in male *ob/ob* mice supplemented or not with oral Gln and subjected to a CRS. Lucifer yellow **(A),** alexa Fluor 680 **(B)** (both measured on colonic using chamber) and plasma FITC **(C)** (Measured after 3h oral gavage) in male *ob/ob* mice (n=9-10/group). Protein expression of occludin **(D)** and mRNA expression of *Ocln* (E), *Cldn4* (F), *Tjp1* (G), *Tjp3* (H), *Cldn15* (I), *Fabp2* (J), *Tjp2* (K), *Cldn12* (L), *Cgn* (M), *F11r* (N) and *Marveld2* (O) in colonic tissue. Groups were compared with 2-way ANOVA followed by Tukey post hoc tests (*p < 0.05, **p < 0.01). Data are presented as mean ± standard error of the mean (SEM).

**Fig. 4.**
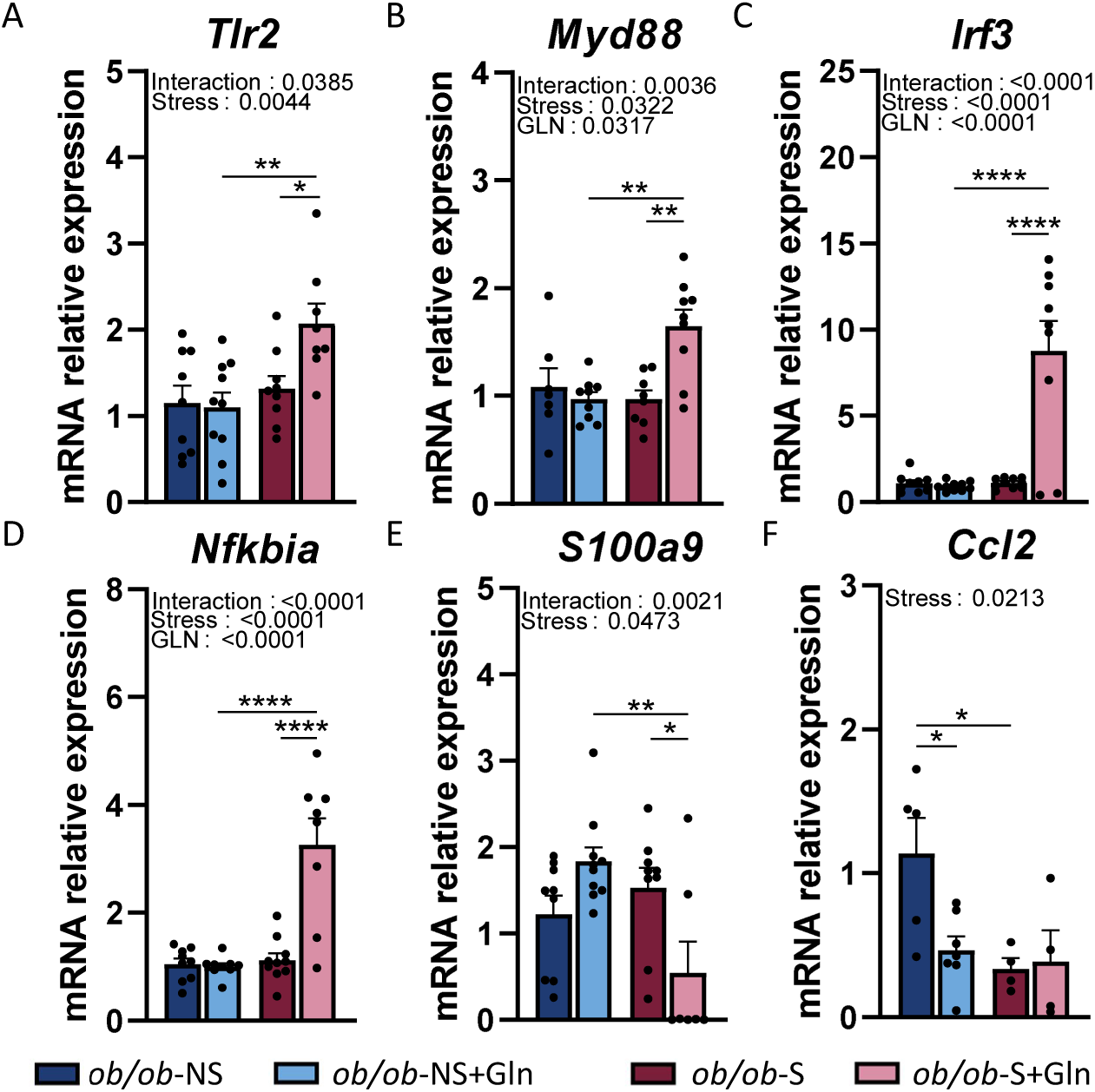
Colonic Inflammation markers in male *ob/ob* mice supplemented or not with oral Gln and subjected to a CRS. mRNA expression of *Tlr2* (A), *Myd88* (B), *Irf3* (C), *Nfkbia*(D), *S100a9* (E) and *Ccl2* (F) (All normalize on *Gapdh* and *Actb*) in colonic mucosa of male *ob/ob* mice (n=9-10/group). Groups were compared with 2-way ANOVA followed by Tukey post hoc tests (*p < 0.05, **p < 0.01, ***p < 0.001, ****p < 0.0001). Data are presented as mean ± standard error of the mean (SEM).

Both stress and Gln, each alone, did not affect the colonic mRNA levels for inflammatory markers (**Fig. 4**), except for the mRNA expression of *Ccl2* that was reduced by both stress and Gln (**Fig. 4F**). By contrast, Gln supplementation in stressed *ob/ob* mice increased the mRNA levels of 4 inflammatory markers (*Tlr2*, *Myd88*, *Irf3*, and *Nfkbia*; **Fig. 4 A-D**) but decreased the mRNA expression of *S100a9* (**Fig. 4E**). No difference was detected for ZO-1 and Claudin-1 protein expression and for mRNA levels for *Cldn2, Cldn3, Cldn8*, *Ticam1*, *Tlr4*, *Tgfb* and *Cxcr3* (**Fig. Supp. 3A-I**).

Taken together, these data suggested that, in *ob/ob* mice, stress and Gln supplementation, each alone, do not alter gut permeability although alterations in colonic tight junction protein expression were observed, despite an increased expression of markers involved in barrier reinforcement.

### Cecal microbial composition in response to stress and Gln supplementation

Cecal microbial community analysis was performed by sequencing the V3–V4 region of the 16S rRNA gene using the Illumina MiSeq platform. We first examined the gut microbiota diversity across the four groups of *ob/ob*-mice. Neither stress nor Gln, when considered independently, affected alpha-diversity as assessed by the Observed species and Chao1 richness indices or by Shannon diversity index. In contrast, Gln supplementation in *ob/ob*-S mice was associated with an increase in richness (Observed species and Chao1) but not in diversity (Shannon index) (**Fig. 5A**). Principal coordinates analysis (PCoA) of Bray–Curtis compositional dissimilarity between cecal samples revealed a significant separation between the communities in response to stress compared to *ob/ob*-NS mice (p<0.05) (**Fig. 5B**). However, Gln supplementation did not significantly alter community structure.

**Fig. 5.**
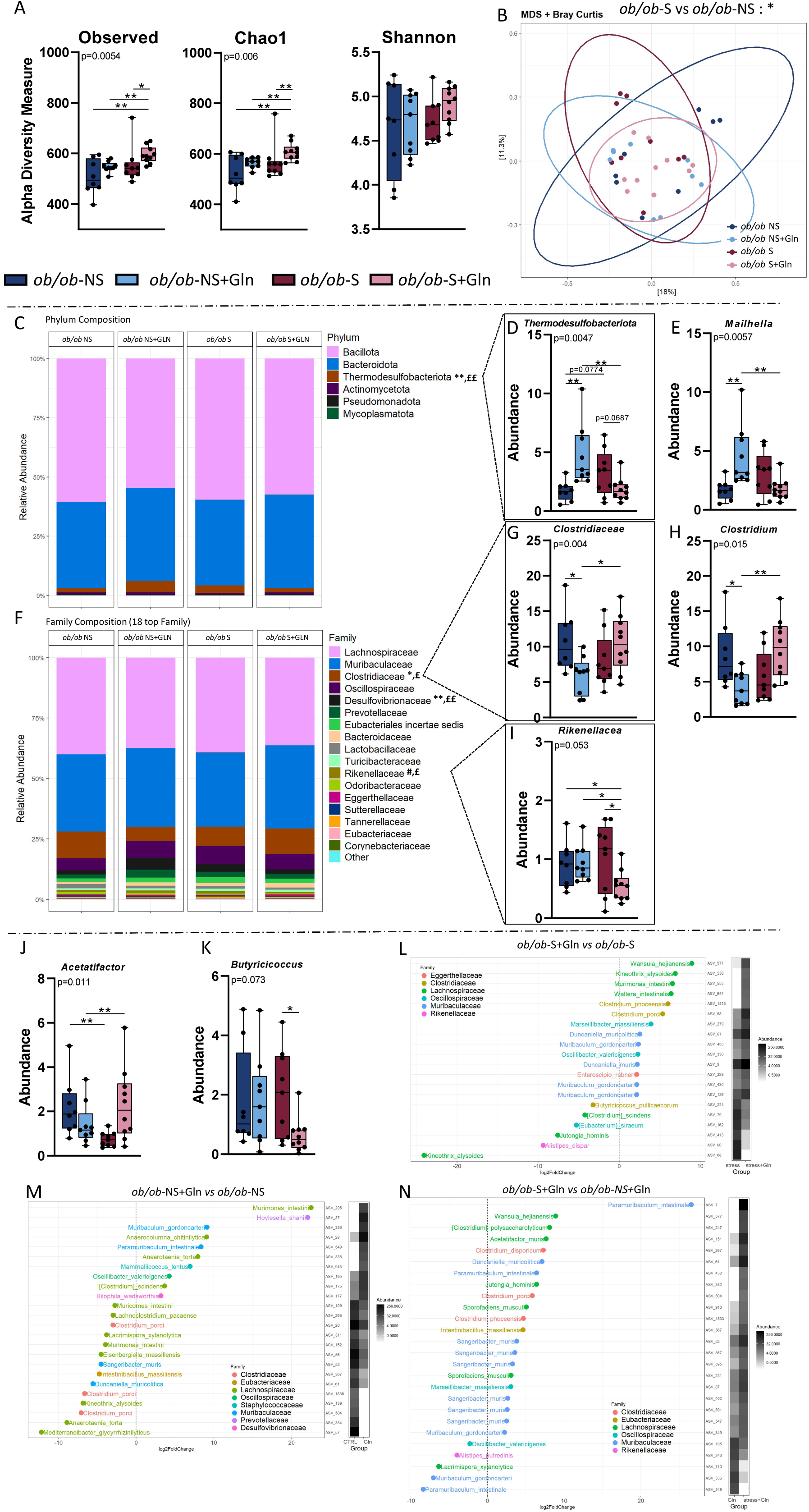
Diversity and Cecal microbiota composition based on 16S rDNA sequencing and graphic representation of differentially abundant ASVs. Observed species richness and Cecal microbial community analysis was performed by sequencing the V3–V4 region of the 16S rRNA gene using the Illumina MiSeq platform (n = 9-10/group). Alpha-diversity indices (Observed species, Chao1 andShannon) are reported in (**A**). Community structure analysis based on Bray Curtis dissimilarity is illustrated by principal coordinates analysis (PCoA) in (**B**), with each dot representing one mouse. Relative bacterial composition at the phylum level (**C**) *Thermodesulfobacteriota* phylum **(D)**, *Mailhella* (**E**) Gln are shown. Relative bacterial composition at the family level (**F**) *Clostridiaceae* family (G), *Clostridium* (**H**) and *Rikenellaceae* (I). Genus *Acetifactor* (J), *butyricicoccus* and differentially abundant ASVs identified in *ob/ob*-S+Gln vs *ob/ob*-NS+Gln (L) *ob/ob*-NS+Gln vs *ob/ob*-NS are displayed in (**M**), while those detected in *ob/ob*-S+Gln vs *ob/ob*-NS+Gln are shown in (**N**). For α-diversity, data are expressed as mean ± SEM and analysed using the Kruskal Wallis test followed by Dunn’s multiple comparisons test (*p < 0.05, **p < 0.01, ****p < 0.0001). Relative abundance data were analysed using Mann–Whitney tests. Differential abundance analysis was performed using log2 fold change (lfc.threshold > 2, min.abundance = 30, adjusted p-value < 0.05). Symbols indicate significant differences between groups: * *ob/ob*-NS *vs ob/ob*-NS+Gln; # *ob/ob*-S *vs ob/ob*-S+Gln; £ *ob/ob*-NS+Gln *vs ob/ob*-S+Gln.

To broadly visualize the gut microbiome composition across groups, we examined the distribution of bacterial phyla and families. In *ob/ob-*NS mice, Gln supplementation increased the relative abundance of the phylum *Thermodesulfobacteriota*, the family *Desulfovibrionaceae* and the genus *Mailhella* (**Fig. 5C–E**), whereas the relative abundance of the family *Clostridiaceae* and the genus *Clostridium* was reduced (**Fig. 5F–H)**. Although stress did not affect significantly these abundances, the effect of Gln supplementation on *Thermodesulfobacteriota* and *Clostridiaceae* observed under unstress condition was no longer apparent in mice exposed to CRS (**Fig. 5D,E,G,H**). In addition, in stress condition, Gln supplementation decreased *Rikenellaceae* abundance compared to other groups (p<0.05) (**Fig. 5I**). Stress reduced the relative abundance of *Acetifactor* in *ob/ob* mice (p<0.01) that was restored following Gln supplementation (**Fig. 5J**). Finally, the abundance of *Butyricicoccus* was decreased in *ob/ob*-S+Gln mice compared to *ob/ob*-S mice (**Fig. 5K**). In *ob/ob*-S mice, Gln supplementation significantly altered the abundance of 11 ASVs, including seven belonging to the *Lachnospiraceae* family (four increased and three decreased) and four belonging to the *Muribaculaceae* family (all increased) (**Fig. 5L**). In *ob/ob*-NS mice, Gln supplementation differentially affected 12 ASVs all belonging to the *Lachnospiraceae* family. Among these, four ASVs were increased and eight were decreased (**Fig. 5M**). Finally, comparison of *ob/ob*-S+Gln and *ob/ob*-NS+ Gln mice identified six differentially abundant *Lachnospiraceae* ASVs that were enriched and one depleted in *ob/ob*-S+Gln mice (**Fig. 5N)**.

The changes in the composition of the bacterial community were associated with alterations in the metabolic activity of the gut microbiota. Indeed, Gln supplementation significantly decreased the total cecal SCFA content including acetate (40% reduction), propionate (40% reduction) and butyrate (67% reduction) while BCFA levels increased (**Fig. 6A-D).**

**Fig. 6.**
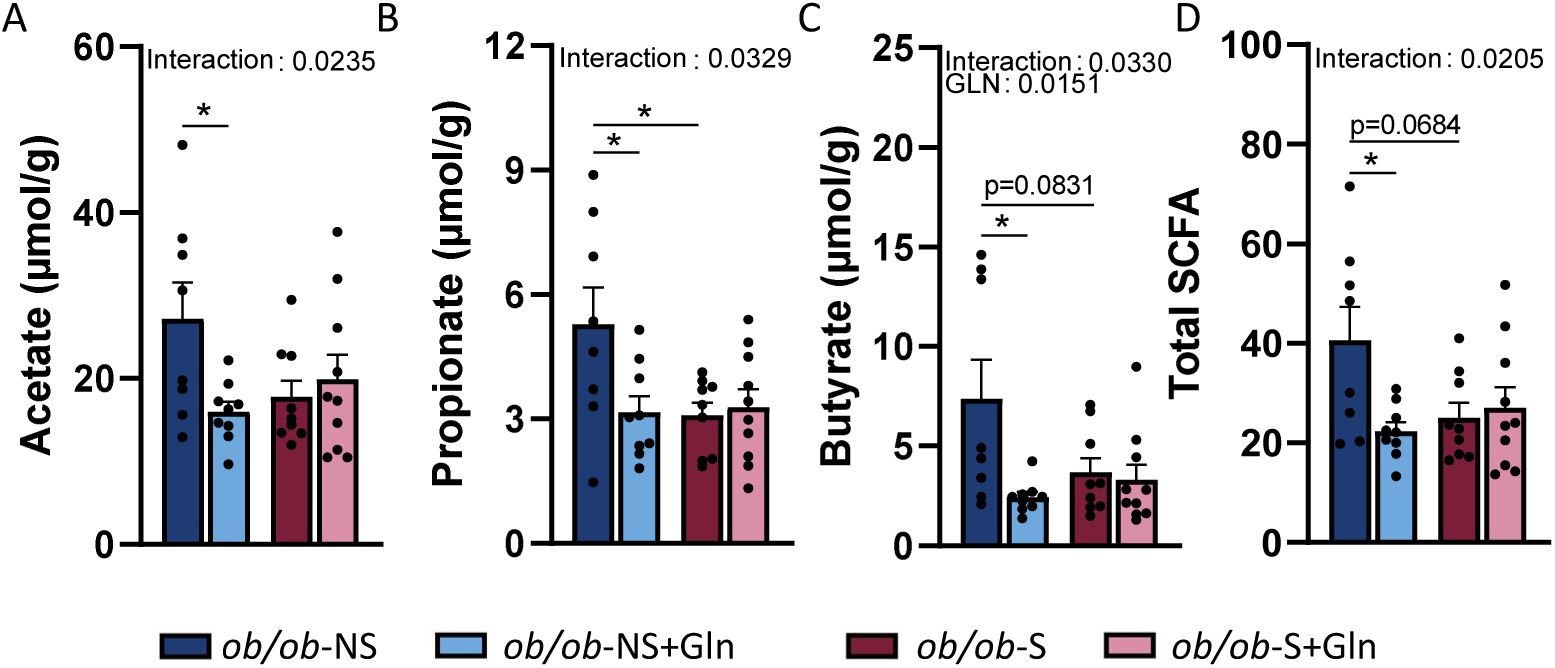
Short-chain fatty acids (SCFA) and branched chain fatty acid (BCFA) in cecal content of male *ob/ob* mice supplemented or not with oral Gln and subjected to a CRS. Changes in microbial metabolic activity were assessed by gas chromatography (n = 9– 10/group). Absolute cecal concentrations (µmol/g) of acetate (**A**), propionate (**B**), butyrate (**C**), total SCFAs (**D)** are shown. Groups were compared with 2-way ANOVA followed by Tukey post hoc tests (*p < 0.05, **p < 0.01, ***p < 0.001, ****p < 0.0001). Data are presented as mean ± standard error of the mean (SEM).

### Correlation analysis between bacterial taxa and cecal SCFAs

In *ob/ob* mice, correlation analysis between cecal bacterial taxa and SCFAs revealed specific associations relevant to obesity. Notably, butyrate showed negative correlations with the family *Staphylococcaceae* (**Fig. 7**). Acetate levels were negatively correlated with *Sutterellaceae* (**Fig. 7**). Propionate levels showed modest correlations, with negative associations with *Oscillospiraceae* (**Fig. 7**). Among the BCFAs, isobutyrate showed the strongest associations with the abundance of bacterial families. Indeed, positive correlations between isobuyrate levels and the abundances in *Staphylococcaceae*, *Atopobiaceae*, *Aerococcaceae* and *Desulfovibrionaceae* were observed, as well as a negative correlation with *Corynebacteriaceae* (**Fig. 7**). Interestingly, the positive correlation with *Atopobiaceae* also existed with valerate and isovalerate levels (**Fig. 7**).

**Fig. 7.**
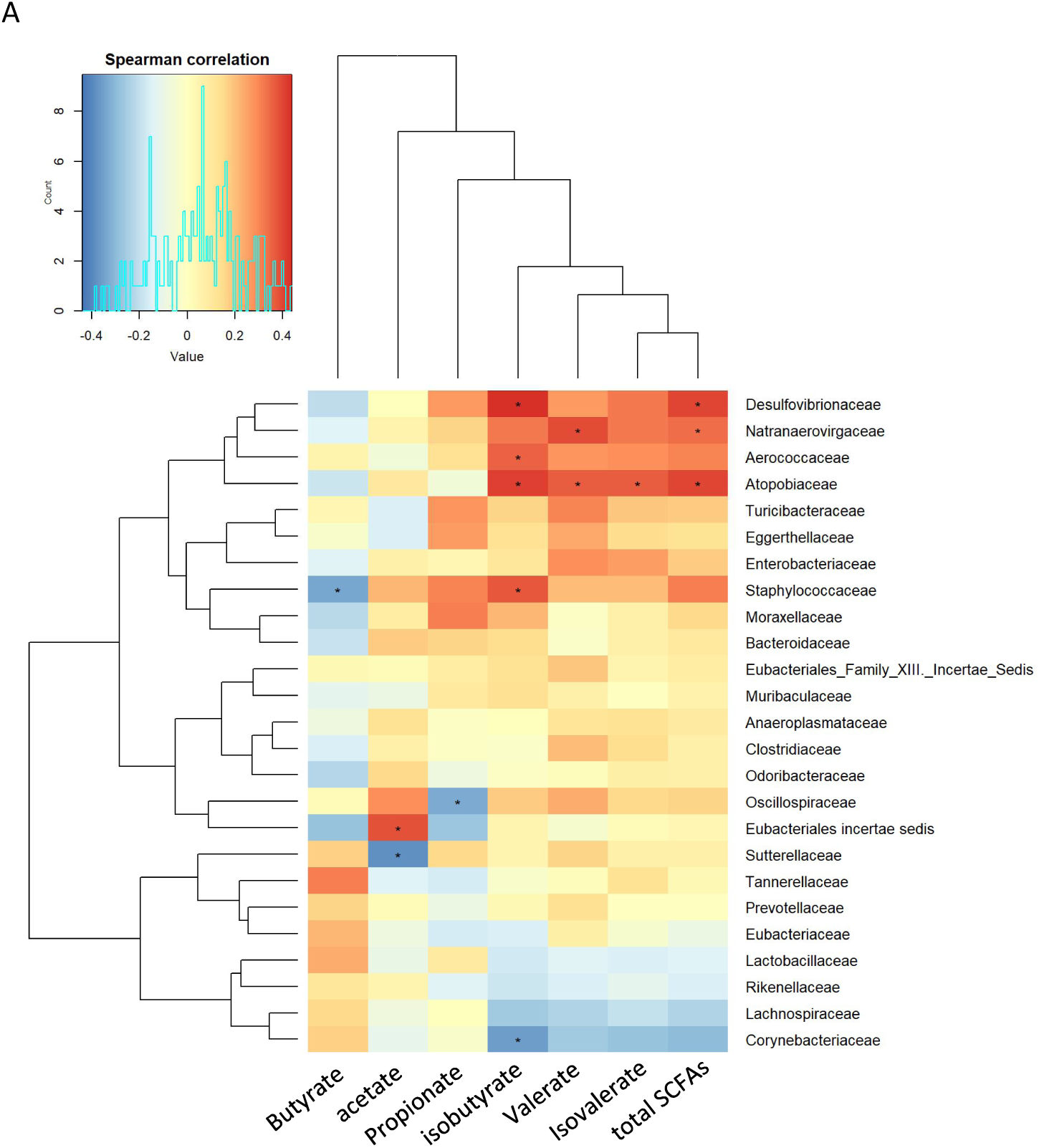
Spearman correlation heatmap between the relative abundance of gut microbiome at the family (A) or genus (B) level and cecal SCFA concentrations in male *ob/ob* mice. Heatmap values are shaded from red (> 0) to blue (< 0) according to correlation scores. The correlation scores have been calculated by using R software version 4.3.3 and heatmaps were created with Pheatmap software version 1.0.12.

## Discussion

Our study investigates the impact of CRS and/or Gln supplementation on metabolic, inflammatory, intestinal and microbial parameters in male obese *ob/ob* mice. Overall, our findings suggest beneficial effects of Gln supplementation in unstressed obese mice but not in stress conditions.

Indeed, Gln supplementation is associated with changes in body composition, insulin resistance, gut microbiota and perigonadal adipose tissue inflammatory response in unstressed *ob/ob* mice. First, in our study, Gln supplementation reduced the gain of body weight, fat and lean mass in unstressed *ob/ob* mice suggesting that Gln may affect energy balance. Interestingly, a reduction in food intake in *ob/ob* mice after an oral Gln supplementation in drinking water with 4% Gln for 40 days has been previously shown, even though that body weight changes were not significant (27). In HFD models, reductions in weight gain after Gln supplementation are not consistently observed since some studies reported a decrease in body weight (37), while others did not (16,30,38). This discrepancy may be related to the duration of Gln supply or to the form of administered Gln: free form or as a L-Alanyl-L-Gln dipeptide. In humans, a pilot study in female patients with obesity showed a weight loss after a 4-week supplementation with Gln (0.5 g/kg body weight/day) (29). Oral Gln supplementation also induced a decrease in fat mass, waist circumference and insulinemia in patients with obesity (39), as well as an increase in postprandial energy expenditure and fat oxidation (40). However, a shorter Gln supplementation period (14 days) was not associated with body weight changes (28,41), underlying the importance of the duration of Gln supplementation. Second, we also observed that Gln supplementation was associated with a reduction in O-GlcNAcylation proteins in the perigonadal adipose tissue, but not in the subcutaneous adipose tissue, in unstressed *ob/ob* mice. Gln supplementation induced a trend for a limitation of inflammatory response specifically in the perigonadal adipose tissue (lower *Tnfa* and *Ccl2* mRNA levels), as well as a reduction in a Gln transporter, *Slc1a5,* mRNA expression. Interestingly, reduced Gln availability in the abdominal subcutaneous adipose tissue of obese individuals has been reported to shift cellular metabolism from glutaminolysis towards glycolysis leading to increased nuclear O-GlcNAcylation and activated inflammatory pathways (42). In our study, Gln supplementation may thus reduce perigonadal O-GlcNAcylation-regulated inflammatory response through an increase of Gln availability. Interestingly, this metabolic benefit was accompanied by a downregulation of the Gln transporter *Slc1a5* in the perigonadal adipose tissue of our *ob/ob* mice. Slc1a5 is a critical driver of cell growth and proliferation, as it provides the Gln necessary for mTORC1 activation (43–45). In the context of obesity, adipose tissue often exhibits an inflammatory, hyper-metabolic phenotype characterized by excessive cellular stress and expansion (46). Recent findings suggest that Gln supplementation can mitigate this by shifting adipose-derived stem cells toward a quiescent metabolic state, resembling that of lean models (46). Therefore, the observed reduction in *Slc1a5* expression in our *ob/ob* mice might represent a beneficial metabolic adaptation mechanism. By reducing Gln transport capacity, the tissue may avoid hyper-proliferation and the associated oxidative stress, ultimately contributing to the overall improvement in health status and reduced inflammatory response (43–46). However, we were not able in the present study to measure Gln content in adipose tissue. The specific response of perigonadal adipose tissue to Gln supplementation compared to subcutaneous adipose tissue requires further investigations. Finally, in our study, Gln supplementation alone affected gut microbiota composition. Even if we did not observe changes in gut microbiota α- and β-diversities after Gln supplementation, the abundance of *Thermodesulfobacteriota* (phylum) and *Desulfovibrionaceae* (family) was increased by Gln supplementation, while the abundance of *Clostridium* (phylum) and *Clostridiaceae* (family) was reduced. In addition, 12 ASVs of *Lachnospiraceae* were affected, as well as SCFA and BCFA caecal contents. These changes in gut microbiota were not associated with major modifications of barrier function and inflammatory response in the colon. Gln is a beneficial amino acid that plays an important role in gut microbiota and immunity by reducing intestinal colonisation and bacterial overgrowth or translocation, and increasing the production of secretory immunoglobulin and anti-inflammatory properties (47,48). These effects may contribute to a more favourable intestinal environment to limit metabolic endotoxemia. Previous studies described the influence of Gln on gut microbiota (48,49) and a pilot study in subjects with obesity reported gut microbiota changes after 14-days of Gln supplementation (50). However, in the present study, we did not provide evidence of the causal role of Gln-induced gut microbiota changes on metabolic parameters. Further experiments using transplantation of microbiota from Gln-receiving mice to naïve mice should be done.

All these data suggest that Gln supplementation may have beneficial effects on adipose tissue during obesity. However, we also demonstrate that Gln response involved in physiological processes, colonic inflammation, and barrier function appears to be altered under stress conditions.

Indeed, we recently reported that Gln modifies the response to CRS in diet-induced obese mice (16) but its impact in genetically obese models remained unknown. In stressed *ob/ob* mice, Gln supplementation fails to reduce body weight, change body composition and metabolic parameters and to reduce the inflammatory response in the perigonadal adipose tissue. In addition, in the present study*, ob/ob* mice exposed to stress conditions exhibited an increase in fasting glucose levels following Gln supplementation. By contrast, male C57BL/6 mice fed with HFD exhibited a decrease in fasting glucose under a similar stress protocol, suggesting a specific effect of Gln in genetically obese models under stress (16). The impact of Gln supplementation on the bacterial abundance and SCFA content observed in unstressed *ob/ob* mice also disappeared, while an increased mRNA expression of colonic inflammatory markers (*Tlr2, Myd88, Irf3, Nfkia*) was observed. The stress and Gln supplementation induced in *ob/ob* mice an increased abundance of sulfate–reducing bacteria, such as those belonging to the *Desulfovibrionaceae*, which have been associated with barrier disruption and pro–inflammatory signalling. It may reflect a shift towards a more dysbiotic gut environment (51,52). However, these bacteria can also have beneficial effects at physiological concentrations. Experimental studies show that H₂S donors can reduce inflammation, enhance mucus production, and help reconstitute the intestinal microbiota biofilm in models of colitis, suggesting a role in mucosal defence and tissue repair rather than toxicity alone. Moreover, H₂S produced by gut microbes is recognized as a regulatory molecule in the gastrointestinal tract, with effects that depend on its concentration and the overall microbial and immune context (53–55). *Acetatifactor* is recognized as a producer of acetate as its main fermentation product in anaerobic conditions, alongside minor butyrate producer (56). Interestingly, Gln supplementation in stressed *ob/ob* mice restored the abundance of *Acetatifactor* and reduced the abundance of *Butyricicoccus* that is traditionally considered to be a butyrate producer, reflecting putative changes in the function of microbial metabolism, even if the fact that taxonomic presence alone does not guarantee significant metabolite output (57,58).

In conclusion, our study reveals that Gln supplementation improves body composition, insulin resistance and inflammatory response in the perigonadal adipose tissue in male genetically obese mice. By contrast, in stress conditions, Gln fails to affect these parameters and increases fasting glucose levels, as well as colonic inflammatory markers. These data are associated with changes in the gut microbiota that deserve further investigations to decipher the underlying mechanisms.

## Acknowledgements

We thank Pamela Lecras and Dr David Vaudry from the PRIMACEN platform (HeRaCleS Inserm US51, CNRS UAR 2026, Université de Rouen Normandie, France) and the staff of GenoToul platform (Toulouse, France).

## Author contributions

Conceptualization, AG, AT, PD, VDo and MC; AT, CL, MM, AG and MC, formal analysis; AT, CE-B, CL, CBF, CG, EM, MM, VDo, VDr, VDo, and AG, investigation; AT, VDo, AG and MC, original draft preparation; writing review and editing, AG, AT, VDo and MC.

---

**Fig. supp. 1.**
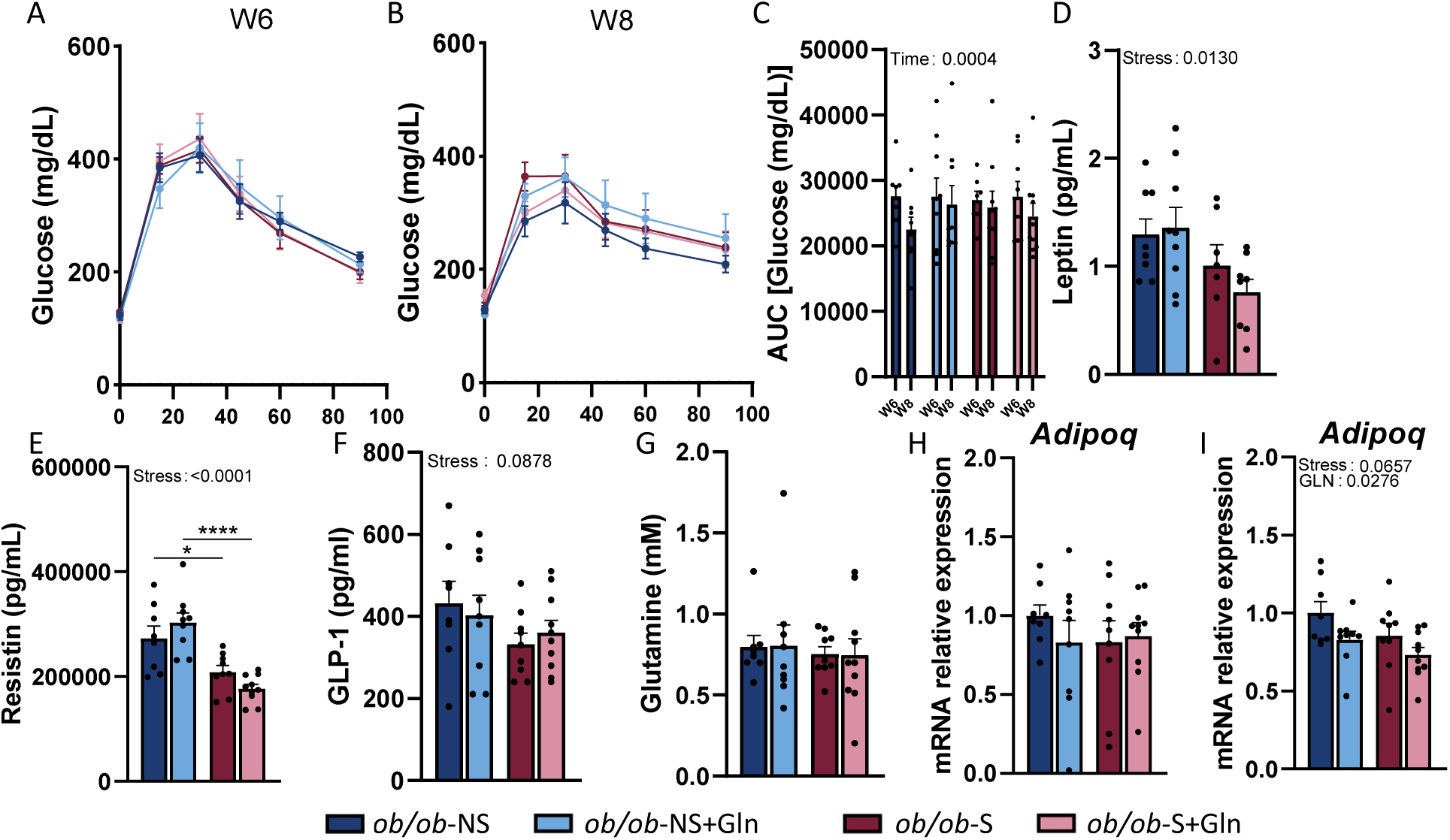
Glycaemic and plasma analysis. Oral glucose tolerance test at W6 **(A),** W8 **(B)** and aera under the curve **(C)** in male *ob/ob* mice. Plasma leptin **(D)** (pg/mL), resistin **(E)** (pg/mL), GLP-1 **(F)** (pg/mL), Gln **(G) (mM),** mRNA relative expression of adiponectin (*Adipoq*) in perigonadal (H) and subcutaneous (I) adipose tissue. Groups were compared with 2-way ANOVA followed by Tukey post hoc tests (*p < 0.05, **p < 0.01, ***p < 0.001). Data are presented as mean ± standard error of the mean (SEM) (n=9-10/group).

**Fig. supp. 2.**
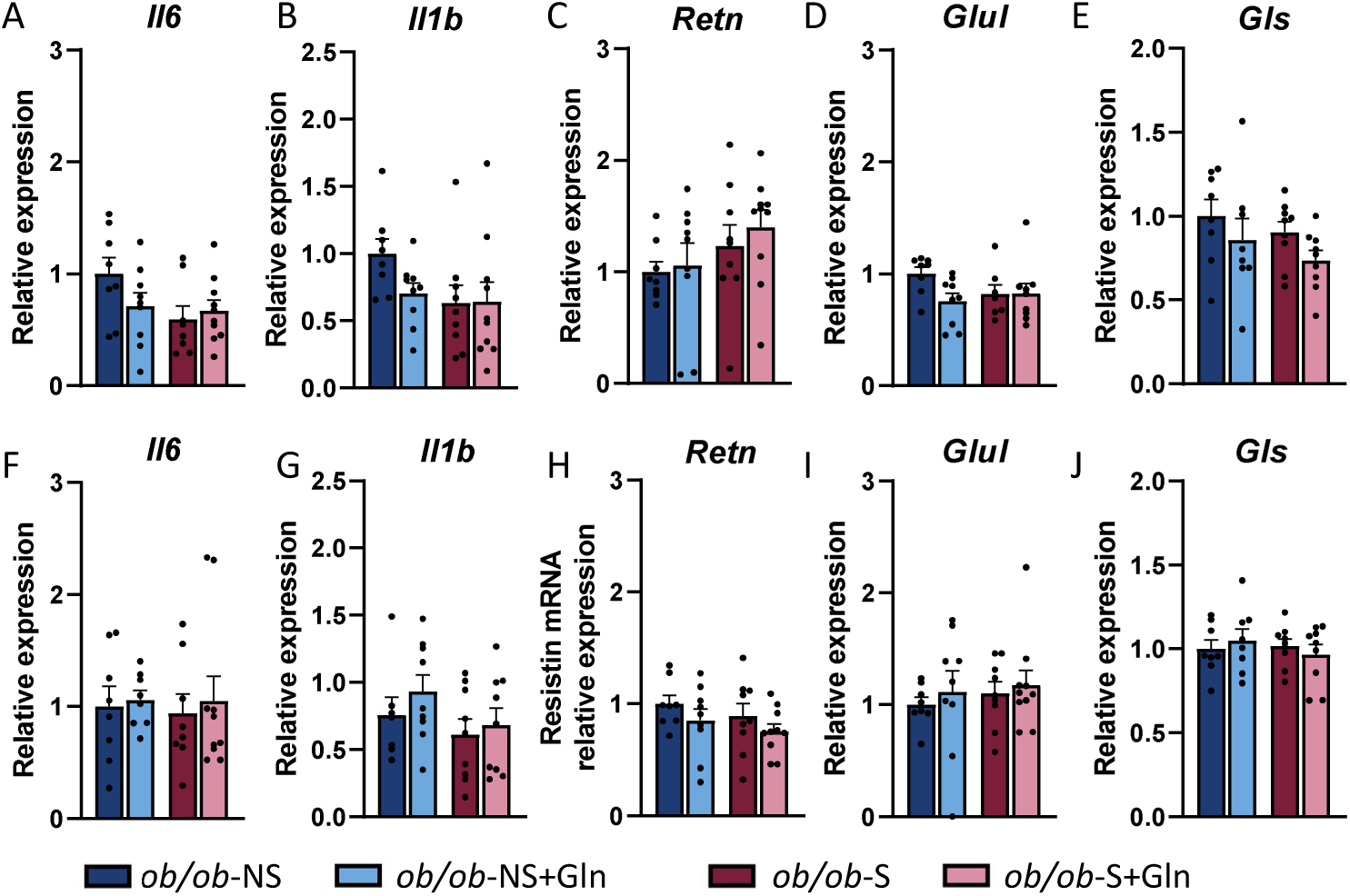
Inflammatory and Gln metabolism markers in perigonadal (A, B, C, D, E) and subcutaneous (F, G, H, I, J) adipose tissues in male *ob/ob* mice. Relative quantification of mRNA transcript levels encoding inflammatory cytokines interleukin 6 (*Il6*; **A, F)**, interleukin 1β (*Il1b*; **B, G)**, resistin (*Retn*; **C, H),** Gln synthetase (*Glul*; **D, I)** and glutaminase (*Gls*; **E, J)** (all normalize on *Rps18* and *B2m)*. Groups were compared with 2-way ANOVA followed by Tukey post hoc tests. Data are presented as mean ± standard error of the mean (SEM) (n=9-10/group).

**Fig. supp. 3.**
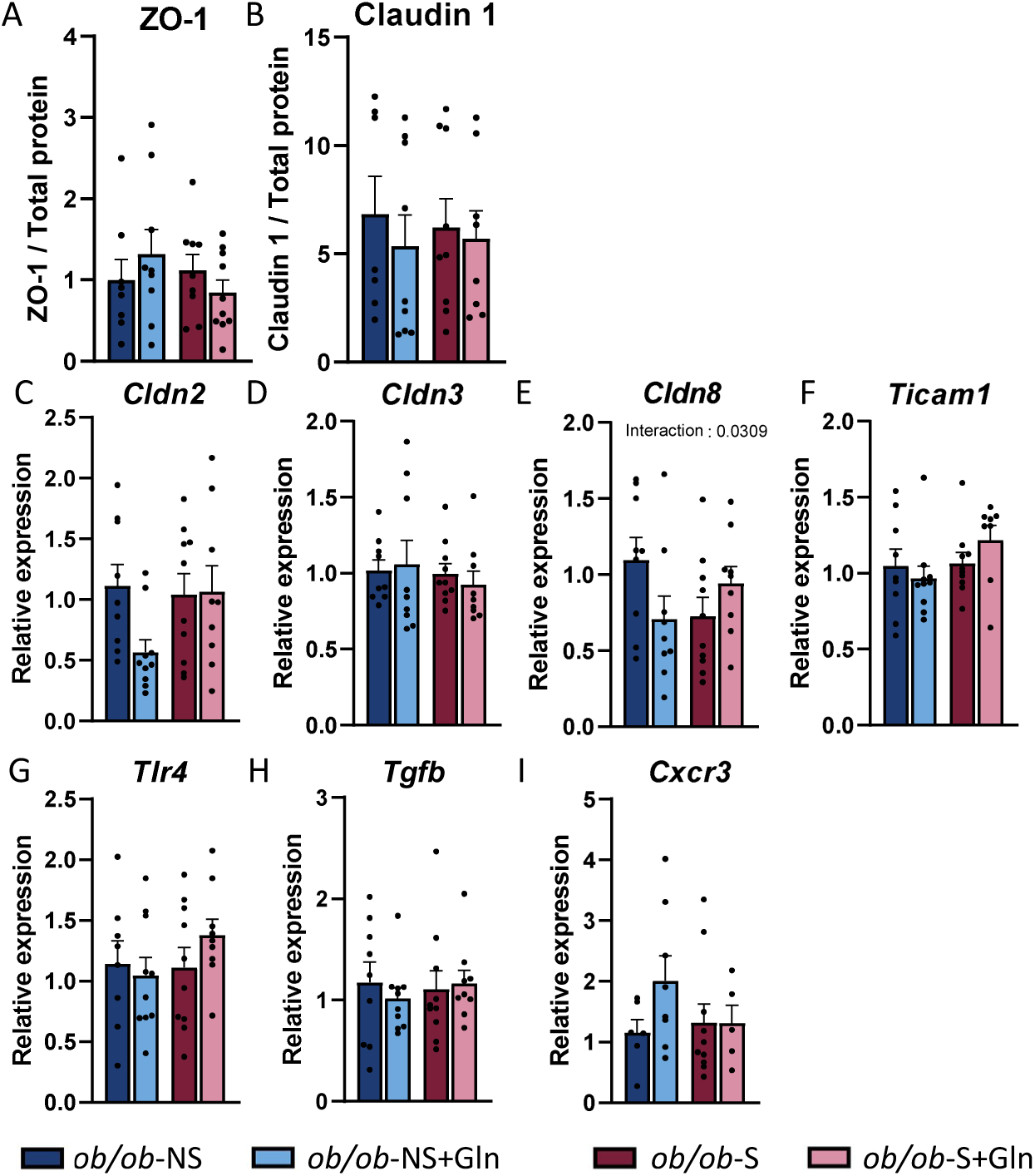
Tight junction protein and inflammation in colonic mucosa of male *ob/ob* mice. Protein expression of Zonula Occludens 1 (ZO-1) **(A)**, Claudin 1 **(B**), and mRNA expression of *Cldn2* **(C)**, *Cldn3* **(D)**, *Cldn8* **(E)**, *Ticam1* **(F)**, *Tlr4* **(G)**, *Tgfb* **(H)**, Cxcr3 **(I)** (All normalize on *Gapdh* and *Actb*) in colonic tissue of males *ob/ob* mice. Groups were compared with 2-way ANOVA followed by Tukey post hoc tests. Data are presented as mean ± standard error of the mean (SEM) (n=9-10/group).

